# Advantages and limits of metagenomic assembly and binning of a giant virus

**DOI:** 10.1101/2020.01.10.902254

**Authors:** Frederik Schulz, Julien Andreani, Rania Francis, Jacques Yaacoub Bou Khalil, Janey Lee, Bernard La Scola, Tanja Woyke

**Author notes:** Contributed equally to the work (Julien Andreani, Rania Francis). Correspondence to: FS, TW.

## Abstract

Giant viruses have large genomes, often within the size range of cellular organisms. This distinguishes them from most other viruses and demands additional effort for the successful recovery of their genomes from environmental sequence data. Here we tested the performance of genome-resolved metagenomics on a recently isolated giant virus, Fadolivirus, by spiking it into an environmental sample from which two other giant viruses were isolated. At high spike-in levels, metagenome assembly and binning led to the successful genomic recovery of Fadolivirus from the sample. A complementary survey of viral hallmark genes indicated the presence of other giant viruses in the sample matrix, but did not detect the two isolated from this sample. Our results indicate that genome-resolved metagenomics is a valid approach for the recovery of near-complete giant virus genomes given that sufficient clonal particles are present. Our data also underline that a vast majority of giant viruses remain currently undetected, even in an era of terabase-scale metagenomics.

## Introduction

Substantial advances in metagenomics and single-cell genomics have rapidly expanded known biodiversity by recovering sequences of 100s of thousands of uncultured bacteria and archaea from the environment and from the human microbiome (Brown et al., 2015; Parks et al., 2017; Rinke et al., 2013; Schulz et al., 2017a). Metagenomics has also recently proven to be a powerful method for assessing diversity and coding potential of environmental viruses (Paez-Espino et al., 2016; Roux et al., 2019). Most viral genomes are small in size and when found in metagenomic data they are readily present on a single contig and thus often considered complete or near complete (Roux et al., 2018). However, this is in stark contrast to genomes of large and giant viruses of the NCLDV, which can be up to several megabases (Abergel et al., 2015; Fischer, 2016). Importantly, recent studies showed that these viruses are among the most diverse and abundant entities in marine systems (Hingamp et al., 2013; Mihara et al., 2018) and are also found in a wide range of non-marine ecosystems (Graham et al., 2019; Kerepesi and Grolmusz, 2017; Roux et al., 2017; Schulz et al., 2018, 2017b; Vavourakis et al., 2016).

Considering the wealth of existing metagenomic data (Chen et al., 2019) it is surprising that comparably few studies describe the recovery of giant virus sequences (Andreani et al., 2018; Bäckström et al., 2019; Roux et al., 2017; Schulz et al., 2018, 2017b; Yau et al., 2011; Zhang et al., 2015). Metagenomic discoveries have preceded the physical isolation of some giant viruses, such as the initial reconstruction of Klosneuvirinae genomes from metagenomic sequences (Schulz et al., 2017b) with subsequent physical isolation of additional members of this proposed viral subfamily, namely Bodo saltans virus (Deeg et al., 2018), Yasminevirus (Hussein Bajrai et al., 2019) and Fadolivirus (Rolland et al., 2019). The genomes from the uncultivated Klosneuvirinae revealed that they encoded comprehensive translation system components (Schulz et al., 2017b), subsequently found in isolated Tupanviruses (Abrahão et al., 2018). Taken together, these studies indicate that metagenomics is of profound value in deriving genomes of giant viruses from the environment, enabling important novel insights into their predicted biology, ecology and evolutionary history. However, this approach is currently lacking validation.

We here conducted a benchmarking experiment to address whether genome-resolved metagenomics provides a valid approach for the recovery of giant virus genomes from environmental sequence data. Spiking viral particles into a wastewater sample, we were able to test the performance of commonly used assembly and binning tools, as well as the ability to detect giant virus genomic information in metagenomes.

## Results

For giant virus co-cultivation experiments, a samples of wastewater was collected from a treatment plant in Toulon France and particles within the sample were sorted by flow cytometry into microplates containing host cells. Co-cultures were monitored by high content screening (see Methods for more details), revealing 10 positive wells on *Acanthamoeba castellanii* strain Neff, while no positive cultures were observed on *Vermamoeba vermiformis*. Giant virus identification by flow cytometry characteristics showed 2 different populations: the first population corresponded to Mimivirus and the second population was unidentified. Scanning electron microscopy showed that 6 wells contained typical Mimivirus-like particles (Figure 1a) and 4 wells contained particles that were 200 to 320 nm in size and resembled Marseillevirus (Figure 1b). The identity of Mimivirus was validated by a specific PCR assay. The genome of the Marseillevirus-like isolate was sequenced and phylogenetic analysis of its DNA polymerase gene confirmed this virus as a new member of the *Marseilleviridae*. We named this virus Phoenician marseillevirus.

**Figure 1.**
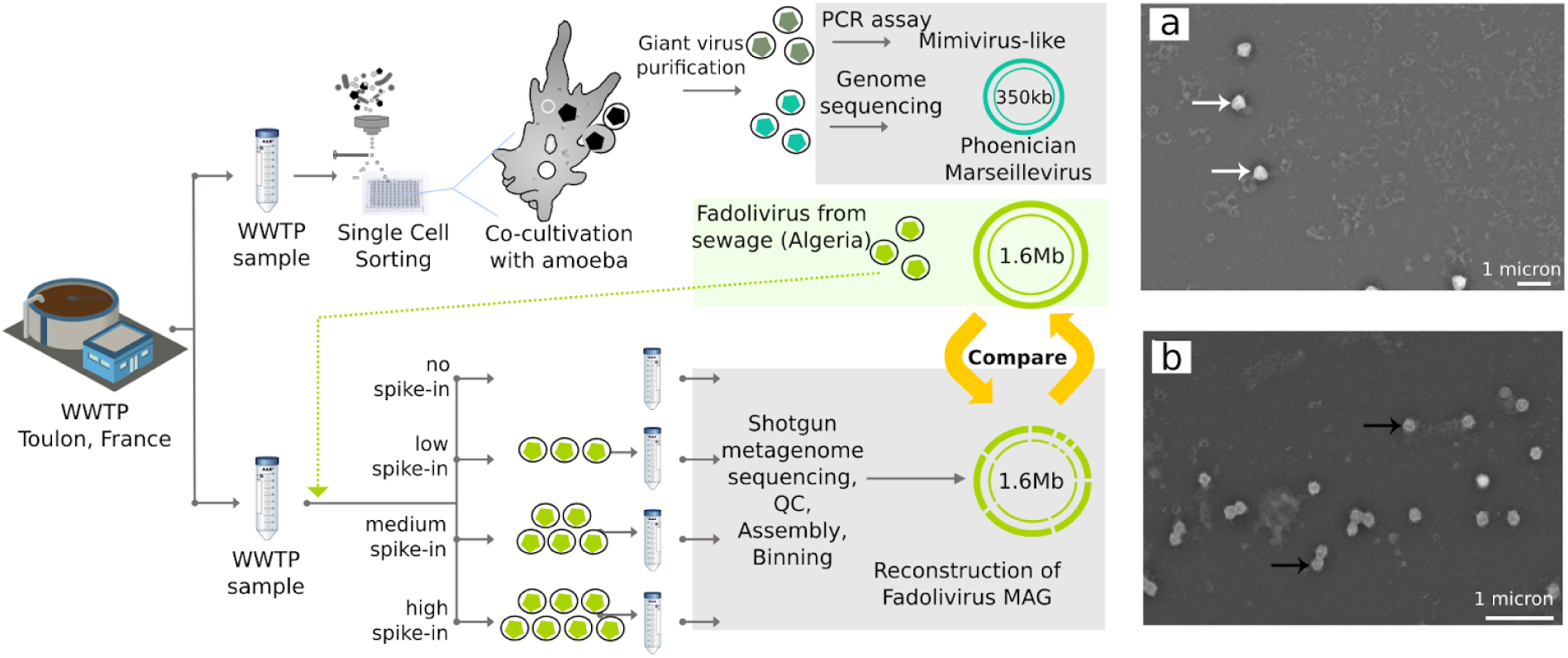
Benchmarking approach to giant virus metagenomics (left panel). Three giant viruses were isolated from wastewater samples by co-cultivation with amoebae; Mimivirus-like particles (dark green), Phoenician marseillevirus (turquoise) and Fadolivirus (light green). Giant virus particles were identified by a specific PCR assay (Mimivirus-like particles) or by whole genome sequencing (Fadolivirus, Phoenician marseillevirus). Fadolivirus particles were purified and spiked in to the initial sample at different concentrations (low, medium, high; see Methods for more details). Samples with and without viral spike-in were subjected to shotgun metagenome sequencing, quality control (QC), assembly and binning. The Fadolivirus metagenome assembled genome (MAG) was then compared to the Fadolivirus reference genome. **Scanning electron micrographs of isolated giant virus obtained with the TM4000 Plus tabletop microscope (right panel). a**. Mimivirus-like particles (white arrows) and **b**. Phoenician marseillevirus particles (black arrows). Scale bars are indicated on each micrograph.

For our metagenomics benchmarking experiment, we began by spiking a portion of the wastewater sample with a known virus, the recently isolated Fadolivirus (Rolland et al., 2019). This viral isolate has a genome size of 1.595 Mb and represents a close relative of Klosneuvirus in the proposed viral subfamily *Klosneuvirinae (Schulz et al., 2017b)*. Samples were spiked with Fadolivirus at the following levels: no (0 viral particles/mL), low (10^3^ viral particles/mL), medium (10^5^ viral particles/mL), or high (10^7^ viral particles/mL); DNA from each sample was sequenced at the DOE Joint Genome Institute. Metagenomics analysis was then performed using a pipeline routinely used for environmental samples, relying on standard QC analysis steps and metaSPAdes (Nurk et al., 2017) assembly (Figure 1). Differential coverage binning with metaBAT 2 (Kang et al., 2019) led to recovery of 114 MAGs. CheckM-based taxonomic classification (Parks et al., 2015) assigned 105 MAGs a bacterial and one MAG an archaeal origin, while 8 MAGs remained unclassified due to the absence of phylogenetic marker genes (Figure 2a). According to MIMAG standards (Bowers et al., 2017), 7 of the MAGs were of high, 44 of medium and 63 of low quality (Figure 2a). The MAG which was predicted to be of archaeal origin (20.3% estimated level of completeness, 4.2% estimated level of contamination; Figure 2a) comprised viral contigs which represented 97.4% of the Fadolivirus reference genome and it did not contain any archaeal sequences.

**Figure 2.**
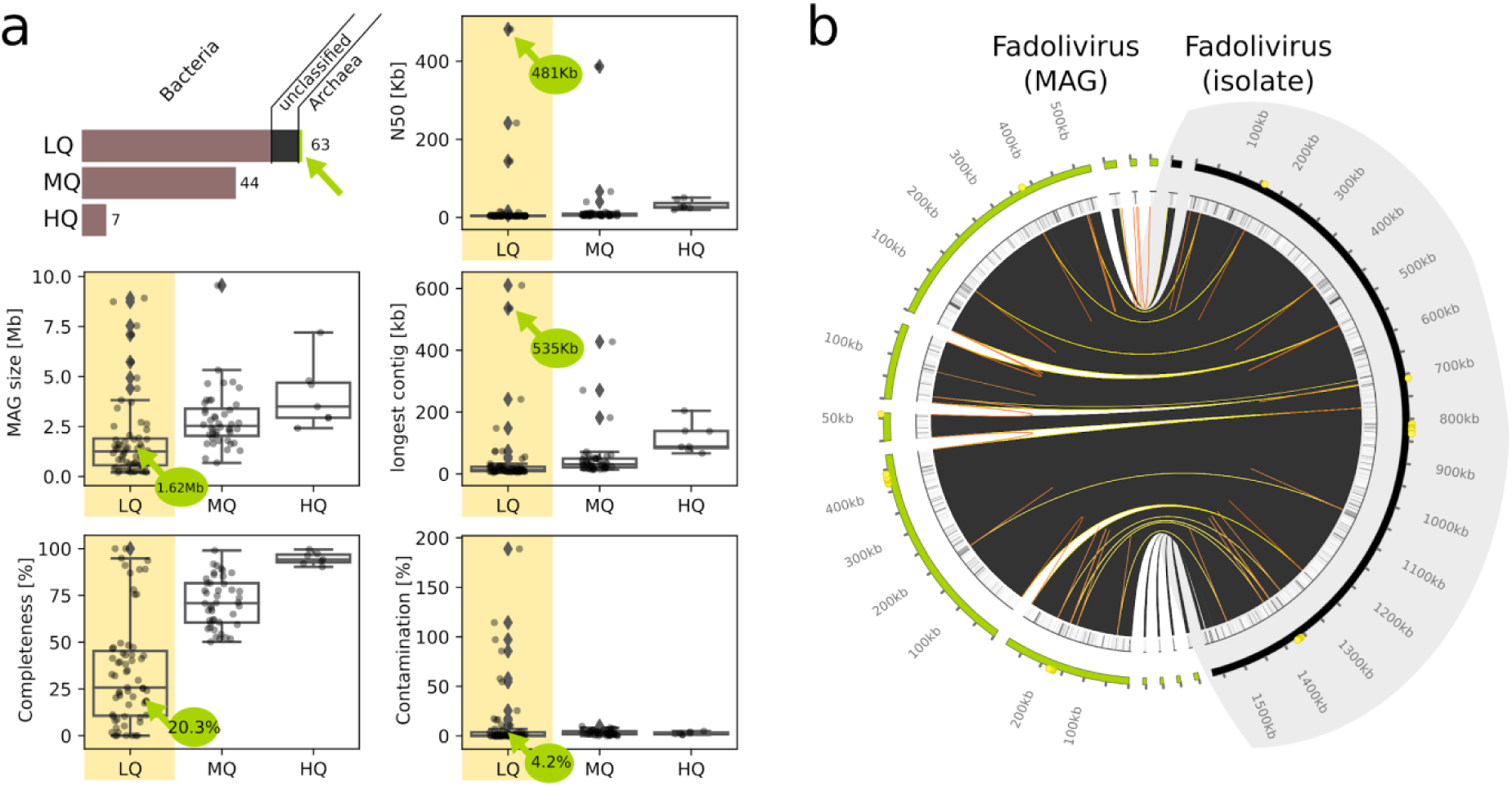
Metagenomic assembly and binning to generate the Fadolivirus metagenome assembled genome (MAG). **a**. Bars indicate the total number of low quality (LQ), medium quality (MQ) and high quality (HQ) MAGs, as defined by MIMAG standards, after differential coverage binning of the metagenome assembly derived from the sample with the highest virus spike-in. Colors indicate domain-level taxonomic assignment of MAGs according to CheckM. Boxplots show different assembly metrics for MAGs. Center lines of box plots represent the median, bounds of boxes the lower and upper quartile, and whiskers extend to points that lie within 1.5 interquartile range of the lower and upper quartile. Green arrows indicate the Fadolivirus MAG. **b**. Whole genome synteny plot of the Fadolivirus MAG (light green) compared to the Fadolivirus reference assembly (black). Areas with > 99% alignment identity between the two assemblies are highlighted in dark grey. For each assembly high identity structural repeats (>95% nucleic acid similarity) with a length of 80-200bp are connected to each other with orange links. Yellow links connect the respective repeats between both assemblies.

This viral MAG was only recovered in the metagenome sample with the high level of viral particle spike-in. Employing differential coverage binning i increased the MAG size to 1.623Mb covering 99.7% of the Fadolivirus reference (Figure 2a, Table 1). The metagenome assembled viral genome had an N50 of 481 kb and comprised 12 contigs, each with a size of at least 5 kb and the largest with a size of 535 kb (Figure 2a, Table 1). Five kb of the Fadolivirus reference genome were missing in the viral MAG. However, the MAG included one additional contig which was not present in the reference genome and two contigs which could only be partially aligned to the reference, totalling 33 kb of unaligned sequence data (Table 1, Figure 2b). Detailed genome comparison of the aligned fraction between the Fadolivirus isolate and MAG did not identify any misassembled regions and revealed 16 mismatches per 100 kb (Table 1). Furthermore, we evaluated the presence of highly identical repetitive sequences within the Fadolivirus genome and found that such sequences were located at the ends of 8 out of 12 contigs of the metagenome assembly (Figure 2b).

**Table 1.** Assembly metrics of the Fadolivirus metagenome assembled genome (MAG) compared to the Fadolivirus reference assembly.

|  |  |
| --- | --- |
| *Reference size [bp] (Fadolivirus)* | 1,595,395 |
| Assembly size [bp] (Fadolivirus MAG) | 1,623,616 |
| Total aligned length [bp] | 1,590,159 |
| Genome fraction [%] | 99.707 |
| N50 [bp] | 481,715 |
| # contigs | 12 |
| Largest contig [bp] | 535,783 |
| # misassemblies | 0 |
| # unaligned contigs | 1 + 2 part |
| Unaligned length [bp] | 33,031 |
| Duplication ratio | 1.026 |
| # N's per 100kb | 0 |
| # mismatches per 100kb | 16.61 |
| # indels per 100kb | 3.52 |

Metagenomic reads from the samples with high, medium, or low levels of Fadolivirus spiked in were mapped to the Fadolivirus reference genome. The high spike-in samples yielded 68-fold more mapped reads than the medium spike-in, and 4,194-fold more mapped reads than the low spike-in (Figure 3a). Metagenomes from the original samples, i.e., those without Fadolivirus spiked in, did not produce any reads that mapped to the genomes of Fadolivirus or Phoenician Marseillevirus. From all metagenomes, sequences of the major capsid proteins (MCPs) were extracted and then compared to sequences of MCPs found in the Fadolivirus and Phoenician Marseillevirus reference genomes and to MCPs available in the NCBI nr database to assign taxonomic origins. Surprisingly, each sample only had between one and six MCPs, which were all on short contigs with low read coverage (Figure 3b). These MCPs showed only low sequence similarity to MCPs of known nucleocytoplasmic large DNA viruses. Fadolivirus MCPs were only detected in samples with the high and medium level of Fadolivirus spike-in and Phoenician Marseillevirus MCPs were not detected in any sample. In the metagenome from the sample with the highest level of Fadolivirus spike-in all Fadolivirus MCP genes were correctly assembled and binned, whereas the samples with the medium level of Fadolivirus spike-in, Fadolivirus MCP genes were present as short fragments distributed over 12 contigs in the unbinned fraction of the metagenome (Figure 3b). We also identified another NCLDV hallmark gene in the metagenomic data, the viral ATPases (NCVOG0249). Phylogenetic analysis showed that the Fadolivirus NCVOG0249 is located on a single contig in the high spike-in sample and split on two contigs in the medium spike-in sample (Figure 3c). We did not recover NCVOG0249 of the Phoenician Marseillevirus, however, we found several ATPases related to Pithoviruses, Kaumoebavirus, Pacmanvirus and others which could not be assigned to known NCDLV lineages (Figure 3c).

**Figure 3.**
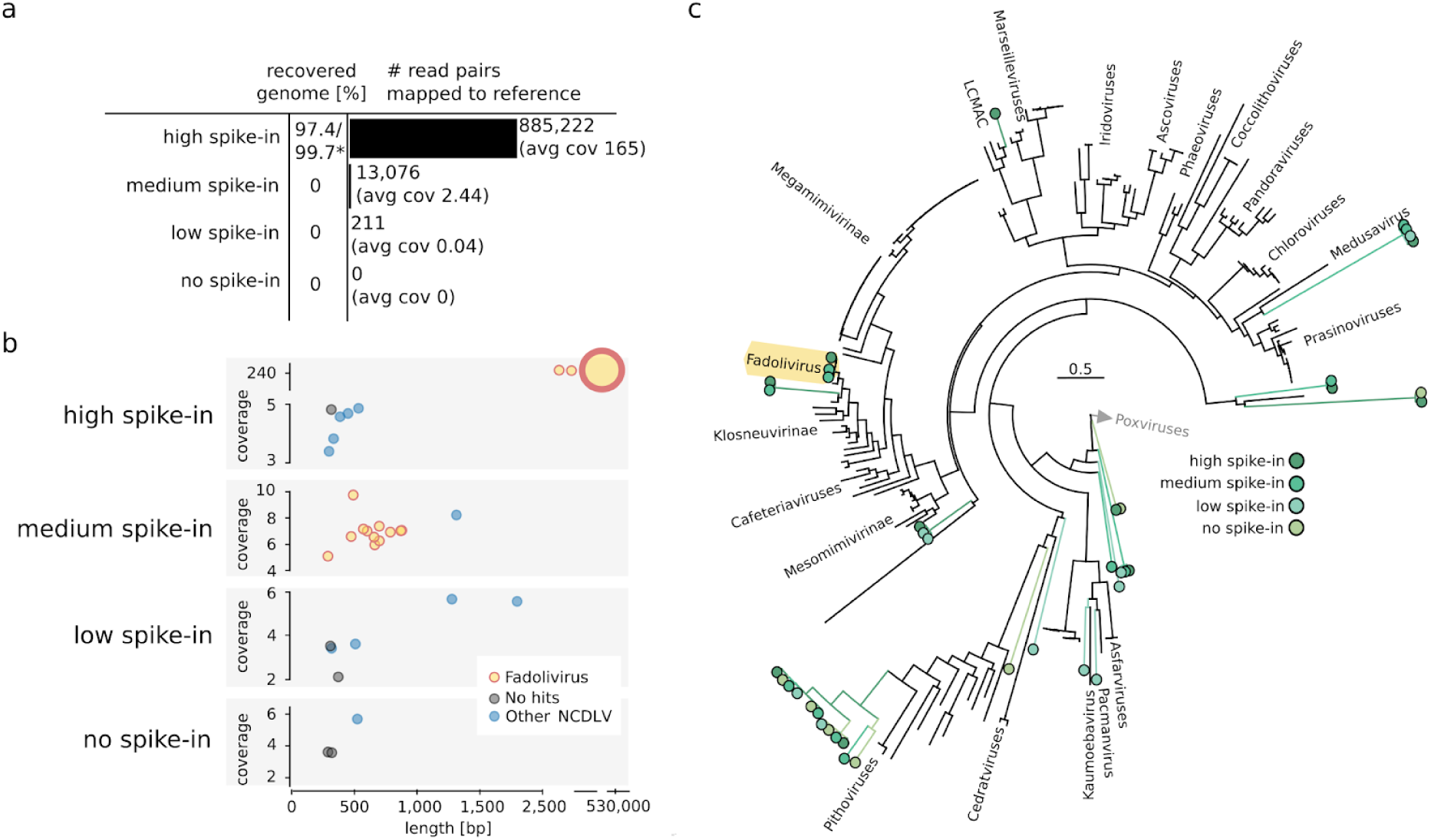
Detection of giant viruses in metagenomic data. **a**. 97.4% and 99.7% of the Fadolivirus genome could be reconstructed in the metagenome with the highest virus spike-in using metaBAT2 (Kang et al., 2019) and differential coverage binning, respectively. Mapping of metagenomic reads from samples with and without viral spike-in to the Fadolivirus and Phoenician marseillevirus reference genomes. The number of mapped reads to the Fadolivirus genome is indicated. No reads mapped to the Phoenician Marseillevirus genome. **b**. Presence of contigs which contained the giant virus MCP gene in samples with and without viral spike-in. Contigs are shown as filled circles which are colored based on the taxonomic origin of the MCP gene. Circle diameter correlates with the total number of MCP genes present on the respective contig. Each contig contained only one MCP gene with the exception of a single contig in the sample with the high viral spike-in which contained 4 copies of the MCP gene. **c**. Phylogenetic positions of viral ATPases (NCVOG0249) extracted from metagenomic data. Scale bar indicates substitutions per site.

## Discussion

Biological insights inferred from genomes extracted from metagenomes rely on sophisticated computational tools and algorithms designed to work efficiently and accurately on diverse sets of environmental sequence data. While these tools are applied on a daily basis by many biologists to answer ecological and evolutionary questions from uncultivated taxa of interest, benchmarking the results often falls short, with the exception of efforts such as CAMI (Sczyrba et al., 2017) and the use of internal standards, such as spike-ins, in some studies (Satinsky et al., 2013; Venkataraman et al., 2018). This is in part due to the difficulty of performing such evaluations using a controlled experiment in a broadly applicable manner. To evaluate the performance of metagenomic assembly and binning of giant viruses we conducted a benchmark experiment, where we spiked particles of a known giant virus, Fadolivirus, into a wastewater sample. Commonly used assembly and binning tools yielded a MAG which represented 97.4% of the Fadolivirus reference genome. By employing differential coverage binning (Kang et al., 2019), we were able to increase Fadolivirus genome recovery to 99.7% (Figure 2). In contrast to the Fadolivirus reference genome assembly, our metagenomic workflow did not yield a closed genome. The presence of assembly breakpoints at highly conserved 80-200 bp repeats demonstrates the difficulty of using the metaSPAdes assembler (Nurk et al., 2017) to resolve such repeats with shorter NovaSeq reads (2×150 bp, average insert size of 241 bp), compared to the longer reads (2×300 bp, average insert size of 253 bp) used for the reference assembly. The performance of the assembler was likely further reduced by the higher complexity of the wastewater sample, which contained more than 100 additional microbial MAGs. However, compared to the microbial MAGs, the Fadolivirus MAG had the highest N50 and contained the largest contig. The Fadolivirus MAG did not have any misassembled regions and had a low mismatch rate of 16 per 100 kb, which would correspond to an accuracy exceeding 99.98%. This comparably high quality of the metagenomic Fadolivirus assembly is likely due to the genomic homogeneity of the clonal Fadolivirus particles that were spiked in. Although this scenario is unlikely to reflect the average environmental sample, our results nicely demonstrate that metagenomics is a powerful tool to recover the nearly complete genome of a giant virus.

Some important aspects for environmental genomics of NCLDV need to be considered. Despite the high completeness of the Fadolivirus MAG when compared to the viral reference, our pipeline classified it as being of archaeal origin and of low quality. While this may be expected, as we used a workflow which relies on a taxonomic framework established for bacterial and archaeal genomes, it also reveals a potential pitfall of metagenome projects. We assessed contamination, completeness and taxonomy with the commonly used tool CheckM (Parks et al., 2015). Building on the CheckM output, MAG quality was then defined according to the MIMAGs standards (Bowers et al., 2017). The lack of most universal cellular marker genes in giant viruses prevents a correct completeness estimate and resulted in the Fadolivirus classification as a “low quality” MAG. The taxonomic classification as archaea can be explained by the fact that the few marker genes which were present in the Fadolivirus genome were most similar to their eukaryotic homologs. The misclassification arises due to the absence of eukaryotic sequences the CheckM reference database (Parks et al., 2015). Importantly, in giant virus metagenomics, misclassification is a known problem, as giant viruses have been deposited as either part of eukaryotic genomes or as bacteria (Andreani and La Scola, 2018; Sharma et al., 2014). In addition, integration of giant virus genes into host genomes can not be excluded (Gallot-Lavallée and Blanc, 2017). Systematically evaluating the performance of our microbial MAG classification workflow on 230 published genomes of large and giant viruses, we found that 70% of them would have been classified as “Archaea” and all of them as “low quality” (data not shown). Thus, it has to be considered that some novel archaeal MAGs in public databases might in fact be misclassified giant viruses. If MIMAG standards (Bowers et al., 2017) are applied, successfully assembled and binned giant virus MAGs should be recoverable from the “low quality” MAG fraction.

Our results show that while a routinely used metagenomics pipeline would likely misclassify and/or not detect giant viruses, a targeted screening would potentially enable the recovery of nearly complete viral genomes. Quality of the resulting MAGs could then be further assessed according to the Minimum Information of an Uncultivated Viral Genome (MIUViG) recommendations (Roux et al., 2018). However, a sample must have a sufficient abundance of giant virus particles for successful recovery and assembly of MAGs, as illustrated by Fadolivirus, which assembled at 165x sequence coverage, but not at 2.4x (Figure 3a). Viruses may naturally be this abundant and clonal after viral replication in eukaryotic hosts. In contrast, viral population heterogeneity or low abundance would complicate genome recovery. Thus, for the recovery of low abundant viruses from complex environmental samples, isolation of giant viruses by co-cultivation with suitable hosts is a highly effective approach (Khalil et al., 2016b; Pagnier et al., 2013).

Our analysis of the MCP and the viral ATPase shows that surprisingly few sequence traces of giant viruses can be found in the wastewater sample despite up to 18 Gb of sequence data generated for each sample and the ability to co-cultivate two NCLDV in the laboratory. Only up to 6 MCPs from NCLDV (other than Fadolivirus) could be detected in each sample. The analysis of the metagenomic viral ATPases revealed the presence of viruses related to giant viruses which had been isolated from wastewater samples in the past, such as Kaumoebavirus, Pacmanvirus and Pithoviruses (Rolland et al., 2019), yet, all of them remained in the unbinned fraction of the metagenomic data. Our results underscore that extraction of NCLDV from metagenomes, even in an era of terabase-scale next generation sequencing, is limited by many lower abundant viruses being beyond the sequence detection level. This does, however, hint at the presence of a vast novelty of currently undetected giant viruses across Earth’s ecosystems.

## Methods

### Sample preparation

Samples were collected in September 2018 from sewage prior to wastewater treatment in Toulon, France (GPS localization: N 43.119; E 5.904). Approximately 1 L of wastewater was transferred to a sterile bottle and then stored at 4°C for 1 month before performing downstream experiments.

### Giant virus co-cultivation

Thirty ml of the wastewater sample were stained overnight with SYBR™ Green I nucleic acid gel stain (Molecular Probes, Life Technologies, USA). Sample was then processed by flow cytometry for sorting using the BD FACSAria™ Fusion cell sorter cytometer (BD Biosciences). After determining 40 populations, sorting was performed in 96 well microplates as previously described (Khalil et al., 2016a). Co-cultivations were then performed on the sorted samples using *Acanthamoeba castellanii* strain Neff and *Vermamoeba vermiformis* as cell hosts, with 10 microplates for each host. Plates were incubated at 32°C and monitored by high content screening for giant virus detection (Francis et al., 2019).

### Giant virus identification

Wells showing potential infection were processed by flow cytometry and scanning electron microscopy (TM4000 Plus microscope, Hitachi High Technologies, Japan) for presumptive identification as previously described (Francis et al., 2019; Khalil et al., 2016b). Virus identification was further validated by PCR and genome sequencing (Ngounga et al., 2013).

### Giant virus spike-in experiment

In parallel and independently of the sample described above, we isolated a novel virus from an Algerian sewage sample (Rolland et al., 2019)by using the same co-culture procedure with *Vermamoeba vermiformis (*Rolland et al., 2019). This virus was named Fadolivirus and we used its particles to artificially contaminate the sample collected from Toulon, France. The rationale for using this particular virus as spike-in was its genome being (i) large in size at 1.6 Mb and (ii) absent from public databases. The latter was critical, as this experiment was a truly blind study in which the US team did not know the identity of the spike-in so as to minimize bias for genomic analysis. Three concentrations of Fadolivirus were selected for the spike-in experiment as follows: each tube containing 35 mL of the homogenate sample contained either (i) 10^3^ viral particles/mL (low spike-in), 10^5^ viral particles/mL (medium spike-in), and 10^7^ viral particles/mL (high spike-in). Another 35mL tube of the sample served as a no spike control. After this step, the 4 tubes were centrifuged using JA-20 rotor at 43,000 × g for 1 hour and 30 minutes in Beckman coulter® Avanti j-26 xp (Beckman, France). The pellets of the 4 tubes were preserved at −80°C before transport and metagenome sequencing and analysis.

Viral particles were quantified by flow cytometry. Data were acquired using log scales for instrument scatter parameters, side scatter (SSC), associated with DNA content detected by Fluorescein (FITC) parameter after SYBR green staining as previously described (Andreani et al., 2017). Thresholds were adjusted on the SSC parameter and 10000 events per sample were acquired. Acquisition and analysis were performed using “BD FACSDiva Software and FlowJo”. The quantification was performed using counting beads (cytocount “DakoCytomation,” a suspension of concentration-calibrated fluorescent microspheres). Absolute count of the population was obtained using the following equation (Khan et al., 2010): (number of cells counted / number of CytocountTM beads counted) × (CytocountTM concentration; i.e.1054 beads/µl) × dilution factor.

### DNA extraction

Metagenomic DNA from each of the four samples (no spike-in; low spike-in, 10^3^ viral particles/ml; medium spike-in, 10^5^ viral particles/ml; high spike-in, 10^7^ viral particles/ml) was extracted using the DNeasy PowerSoil Kit (Qiagen, Germantown, MD). As the samples were liquid, the manufacturer’s protocol was adjusted as follows: briefly, 35 ml of wastewater samples were centrifuged for 45 min at 10,000 rpm at 4°C. The supernatant was decanted and the resulting pellet resuspended in 500 μl of reserved supernatant. The resuspended pellet was then deposited in the kit’s bead tube, in place of soil. The manufacturer’s protocol was followed thereafter. All DNA extracts were quantified using the PicoGreen assay and the Qubit 2.0 Fluorometer (Invitrogen, Carlsbad, CA).

### Library creation and sequencing

Sequencing libraries were created using the TruSeq DNA PCR-Free DNA Sample Preparation Kit, following the manufacturer’s protocol (Illumina, San Diego, CA). Libraries were sequenced on the Illumina NovaSeq platform (2 × 150 bp) at the U.S. Department of Energy’s Joint Genome Institute (JGI) yielding between 14 and 18 Gb of sequence per sample.

### Metagenome assembly and binning

Reads were corrected using bbcms 38.34 (http://bbtools.jgi.doe.gov) with the following command line options: bbcms.sh metadatafile=counts.metadata.json mincount=2 highcountfraction=0.6 in=out.fastq.gz out=input.corr.fastq.gz. The readset was assembled using metaSPAdes assembler with metaspades 3.13.0 (Nurk et al., 2017). This was run using the following command line options: spades.py -m 2000 --tmp-dir scratch -o spades3 --only-assembler -k 33,55,77,99,127 --meta -t 72 -1 reads1.fasta -2 reads2.fasta.

The input read set was mapped to the final assembly and coverage information was generated with bbmap 38.34 (http://bbtools.jgi.doe.gov). This was run using the following command line options: bbmap.sh nodisk=true interleaved=true ambiguous=random in=out.fastq.gz ref=assembly.contigs.fasta out=pairedMapped.bam covstats=covstats.txt bamscript=to_bam.sh. Genecalling was performed with prodigal (Hyatt et al., 2010) using the -meta option. Contigs were organized into genome bins based on tetranucleotide sequence composition with MetaBat2 (Kang et al., 2019).

### Identification of the Fadolivirus MAG

To make this a blind study preventing any bias in the sequence data processing and analysis, the Fadolivirus reference sequence and any information about this isolate was kept in the LaScola laboratory until the viral MAG data was generated and analyzed at the JGI. The data were then revealed and compared. Diamond blastp was used to compare metagenomic proteins against the Fadolivirus reference genome (Buchfink et al., 2015). Only one MAG contained the proteins found in the Fadolivirus reference genome, which was used for detailed comparison with the Fadolivirus reference genome using QUAST (Mikheenko et al., 2016), nucmer from the MUMMER package (Delcher et al., 2003) and Circos to generate a whole genome synteny plot (Krzywinski et al., 2009).

### Survey of the major capsid protein and the viral ATPase

A set of HMMs for the NCLDV MCP gene and the viral ATPase (NCVOG0249) was used in hmmsearch (version 3.1b2, hmmer.org) with a cutoff of 1e-10 to identify putative MCP genes on metagenomic contigs. Resulting protein hits were extracted from the metagenome and subjected to diamond blastp (Buchfink et al., 2015) against the nr database (May 2019) and against all proteins found in the Fadolivirus reference genome. Extracted NCVOG0249 were then aligned with MAFFT-linsi (v7.294b) (Katoh and Standley, 2016) and the resulting amino acid alignments trimmed with trimal (v1.4, -gt 0.9) (Capella-Gutiérrez et al., 2009). Phylogenetic trees were built using IQ-tree (v1.6.12) (Nguyen et al., 2015) with LG+F+R5 (ATPase) based on the built-in model select feature (Kalyaanamoorthy et al., 2017) and 1000 ultrafast bootstrap replicates (Hoang et al., 2018). The ATPase phylogenetic tree was visualized with iTol (v5) (Letunic and Bork, 2016).

## Acknowledgements

This work was conducted by the US Department of Energy Joint Genome Institute, a DOE Office of Science User Facility, under Contract No. DE-AC02–05CH11231 and made use of resources of the National Energy Research Scientific Computing Center, also supported by the DOE Office of Science under Contract No. DE-AC02–05CH11231.

## Declaration of Competing Interests

The authors declare no competing interests.

